# Distinct optokinetic reflex phenotypes in Frmd7 and Chrnb2 mutant mice

**DOI:** 10.64898/2026.04.03.716267

**Authors:** Jiaqi Qi (戚加奇), Akihiro Mastumoto, Keisuke Yonehara

## Abstract

**Highlights:** Binocular rotational stimulation enhances OKR in wild-type mice along both axes.

Chrnb2^tm^ mice show spontaneous horizontal eye oscillations regardless of input.

Frmd7^tm^ mice lack binocular enhancement of vertical OKR.

The optokinetic reflex (OKR) stabilizes retinal images during global motion and depends on retinal direction-selective (DS) circuits. Although multiple mutant mouse strains exhibit impaired DS circuits via distinct mechanisms, differences in OKR phenotypes remain unexplained. Here, we developed a behavioral system to quantify mouse eye movements under controlled rotational and translational visual stimuli.

Using this platform, we examined OKR in wild-type (WT) mice and two DS circuit mutants: Frmd7^tm^ mice, which lack horizontal DS tuning and horizontal OKR, and Chrnb2^tm^ mice, which have disrupted β2-nAChR-dependent cholinergic spontaneous activity during development. Consistent with previous research, both mutants lacked horizontal OKR across conditions, while vertical OKR was preserved. We found that Chrnb2^tm^ mice exhibited spontaneous horizontal eye oscillations regardless of visual input. This phenotype was absent in Frmd7^tm^ mice, suggesting that defective retinal waves in Chrnb2^tm^ mice may induce circuit-level instability distinct from the loss of horizontal DS tuning alone. In addition, WT mice showed enhanced vertical OKR under binocular rotational stimulation, which was absent in Frmd7^tm^ mice.

Together, these findings provide a functional comparison of Frmd7^tm^ and Chrnb2^tm^ mice and establish a quantitative framework for dissecting how specific genetic perturbations alter the retinal computations underlying horizontal and vertical OKR.

## Introduction

Visual stability is essential for accurate perception of the external world. During natural behaviors, even slight head movements could displace retinal images, resulting in motion blur and reduced spatial resolution(Wurtz, 2008). To counteract this instability, vertebrates have evolved several compensatory eye movement reflexes, among which the OKR plays a central role in stabilizing gaze during sustained visual motion(Stahl, 2004). The OKR is characterized by a slow eye-tracking movements followed by a fast resetting saccades, allowing the eyes to continuously follow the direction of the moving stimuli(van Alphen et al., 2001). In rodents, OKR can be easily evoked by presenting visual gratings to head-fixed animal, making it a widely used assay for probing visual motion processing and eye movement circuits(Douglas et al., 2005).

This behavior relies on the accessory optic system (AOS), a group of brainstem nuclei that receive direct input from retinal direction-selective ganglion cells (DSGCs) (Giolli et al., 2006; Simpson, 1984). Visual motion information is first extracted in the retina by DSGCs, which encode the direction and velocity of global image motion(Dhande & Huberman, 2014; Yonehara et al., 2009). There are two classical types of DSGCs in the mouse retina. ON–OFF DSGCs project primarily to the dorsal lateral geniculate nucleus(Stewart et al., 1971) and superior colliculus(Vaney et al., 1981). In contrast, ON DSGCs project to the nuclei of the AOS(Simpson et al., 1979) and are specifically dedicated to mediating the OKR(Yoshida et al., 2001). Accordingly, horizontal-preferring ON DSGCs project to the nucleus of the optic tract (NOT) and dorsal terminal nucleus (DTN), key components of the accessory optic system, which relay these signals to brainstem oculomotor nuclei to generate compensatory horizontal eye movements.(Dhande et al., 2013; L. O. Sun et al., 2015). In contrast, vertical-preferring ON DSGCs project to the medial terminal nucleus (MTN), a major component of the AOS specialized for encoding upward and downward motion(Dhande et al., 2013; Giolli et al., 2006; L. O. Sun et al., 2015).

From the AOS, motion signals are transmitted to the vestibular complex through the inferior olive and the medial vestibular nucleus (MVN), where visual and vestibular inputs converge to refine estimates of head and image movement(Balaban, 1988; Cullen, 2012; Giolli et al., 2006). The vestibular nuclei then provide monosynaptic and polysynaptic projections to the brainstem oculomotor nuclei—including the abducens nucleus (cranial nerve VI) and the oculomotor nucleus (cranial nerve III)—which contain the motoneurons that directly innervate the extraocular muscles(Highstein & Holstein, 2006; Horn & Leigh, 2011). These final motor pathways generate the slow tracking component of the OKR and participate in coordinating the fast saccadic reset through premotor burst neuron circuits(Cullen, 2012; Horn & Leigh, 2011; Scudder et al., 2002). From retinal DSGCs to AOS, vestibular nuclei, and oculomotor motoneurons, this pathway forms the neural architecture that enables precise stabilization of visual scenes during self-motion or motion of the visual surround(Dhande & Huberman, 2014; Giolli et al., 2006; Cullen, 2012; Highstein & Holstein, 2006; Horn & Leigh, 2011; Scudder et al., 2002).

Within the retina, direction selectivity is generated by asymmetric inhibitory and excitatory inputs from starburst amacrine cells (SACs) to DSGCs (Barlow & Hill, 1963; Taylor & Vaney, 2003; Demb, 2007; Fried et al., 2002; Yonehara et al., 2011; Elstrott et al., 2008). These retinal circuits are highly specialized, with distinct DSGC subtypes tuned for horizontal versus vertical motion(Kay et al., 2011; Mauss et al., 2017; Trenholm & Awatramani, 2015). Disruption of these circuits leads to selective impairment of OKR subtypes. For example, mutations in Frmd7, a gene associated with human congenital nystagmus, specifically abolish horizontal direction selectivity without affecting vertical tuning. In Frmd7^tm^ mice, this deficit results in a complete loss of horizontal OKR, while vertical OKR remains intact(Tarpey et al., 2006; Yonehara et al., 2016). Thus, Frmd7^tm^ mice provide a valuable model for dissecting the retinal mechanisms underlying horizontal OKR tracking.

Another molecule critical for retinal DS circuit development is β2-nicotinic acetylcholine receptors (β2-nAChRs), encoded by the Chrnb2 gene. Chrnb2 is highly expressed in SACs and is required for cholinergic signaling that drives spontaneous neural activity during development, called retinal waves(Ackman et al., 2012; Burbridge et al., 2014; Feller, 1999; Stafford et al., 2009). Chrnb2^tm^ mice display profound defects in retinal direction selectivity; horizontal direction selective tuning is largely reduced than vertical(Burbridge et al., 2014; Tiriac et al., 2022). However, an important difference exists between species; humans carrying the FRMD7 mutation exhibit characteristic horizontal eye oscillations, while such oscillatory eye movement is not observed in Frmd7^tm^ mice(Yonehara et al., 2016). Therefore, it remains unclear whether disruption of the Chrnb2 gene leads to abnormal oscillatory eye movements in mice.

To address this question, we developed a behavioral recording system, which can independently measure horizontal and vertical movements in head fixed mice. By systematically varying stimulus parameters, including optical flow-related motion types (rotational vs. translational), spatial frequency and temporal frequency, we first determined the optimal visual conditions for eliciting robust horizontal and vertical OKR in WT mice. We then used these optimized paradigms to compare eye movement responses across WT, Chrnb2^tm^ and Frmd7^tm^ mice. Consistent with previous findings, Frmd7^tm^ mice completely lacked horizontal OKR while retaining vertical OKR. Importantly, we found that WT mice exhibited enhanced vertical OKR responses under binocular rotational stimulation, whereas this enhancement was absent in Frmd7^tm^ mice. Chrnb2^tm^ mice also failed to generate horizontal OKR under all stimulation conditions tested. Strikingly, Chrnb2^tm^ mice exhibited significant spontaneous horizontal eye oscillations in the absence of visual stimulation, and this oscillation could not be eliminated by any visual stimulation pattern, while no such oscillation was observed in WT or Frmd7^tm^ mice. The demonstrating that Chrnb2 is indispensable for the development of horizontal direction-selective circuits, which is required for horizontal reflex tracking. Furthermore, these results reveal an unexpected link between cholinergic signaling disorders and the occurrence of oscillatory eye movements. Together, these results directly link retinal circuit deficits to their behavioral consequences, providing new insights into the neural mechanisms that support horizontal and vertical OKR.

## Results

To record both horizontal and vertical OKR, we designed an experimental setup to monitor eye movements in head fixed mice (Figure 1A). In this setup, the head fixed mouse was placed on an adjustable-height platform, with two monitors positioned on either side. Moving gratings were presented to elicit OKR in either the horizontal (Figure 1B,C) or vertical direction (Figure 1B,D). Horizontal visual stimulation stimulated head rotation along the yaw axis, whereas vertical visual stimulation simulated rotation along the roll axis. To prevent the camera from obstructing the mouse’s visual field and disrupting the continuous presentation of visual stimuli to the retina, a 45° hot mirror was positioned to the left of the mouse. The movement of the left pupil was recorded by reflecting it into the camera via the hot mirror.

**Figure 1.**
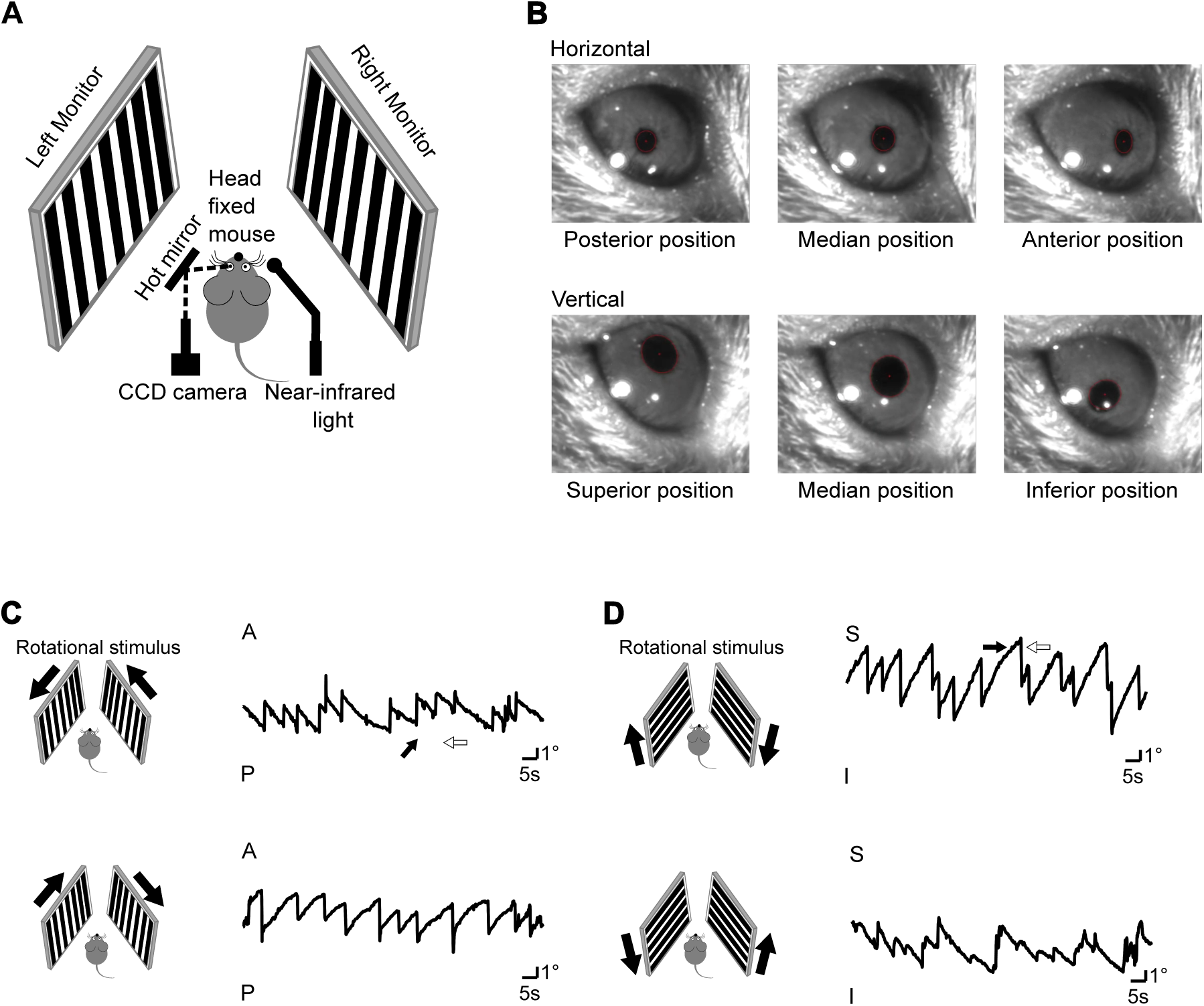
Experimental setup for recording horizontal and vertical OKR. (A) Schematic of the experimental setup. A 45° hot mirror was positioned to the left of the mouse, with a CCD camera placed behind the mouse on the left side and a near-infrared light source positioned behind the mouse on the right side. (B) Pupil position of the left eye during rotational stimulation. Top, three representative pupil positions during horizontal rotational stimulation: slow tracking toward the posterior side (left), resetting position (middle), and slow tracking toward the anterior side (right). Bottom, three representative pupil positions during vertical rotational stimulation: slow tracking toward the superior side (left), resetting position (middle), and slow tracking toward the inferior side (right). (C and D) Changes in pupil position over time during rotational stimulation. Arrows indicate the direction of stimulation. (C) Horizontal direction. Top, pupil position over time during posterior directed stimulation. Bottom, pupil position over time during anterior directed stimulation. (D) Vertical direction. Top, pupil position over time during superior directed stimulation. Bottom, pupil position over time during inferior directed stimulation.

We evaluated the efficiency of OKR elicitation by measuring the average eye tracking movements (ETMs) over one minute. One reflex was defined as a slow tracking movement followed by a resetting saccade, and ETMs was quantified by counting the number of resetting saccades (Methods). Some experimental setups have limitations in reliably eliciting downward (inferior) OKR(Kiraly et al., 2024). Before characterizing the mutant phenotype, we first verified that the setup reliably elicits OKR in four cardinal directions using rotational visual stimuli in WT mice (Figure 1C,D; see also Video S1 and S2).

### Rotational stimulation more effectively elicits OKR than translational stimulation and reveals optimal spatiotemporal frequency tuning

Previous studies have demonstrated that the strength of the OKR is highly influenced by both the type of visual motion and its spatiotemporal characteristics(Sugita et al., 2015; Yoshida et al., 2001). However, most of these investigations have primarily focused on horizontal OKR(Cahill & Nathans, 2008; Tabata et al., 2010), leaving the optimal visual conditions for eliciting vertical OKR less well understood.

To determine the visual parameters that most effectively elicit both horizontal and vertical OKR in head fixed WT mice, we systematically manipulated three key stimulus variables: temporal frequency (TF), spatial frequency (SF), and motion pattern (rotational vs. translational) of the visual field, which simulated either head rotation or translation of the animal. This experimental design enabled a direct comparison of how different visual conditions affect the robustness of OKR responses.

Our results revealed that rotational visual stimulation consistently evoked stronger OKR than translational stimulation when other parameters were held constant (Figure 2A, B). Specifically, the curve of pupil position change over time during the presentation of horizontal visual stimulation shows that, rotational horizontal (RH) stimulation can always evoke more reflexes than translational horizontal (TH) stimulation (Figure 2A). Further, horizontal OKR elicited by RH stimuli exhibited significantly higher ETMs per minute compared to those elicited by TH grating (Figure 2C). Similarly, rotational vertical (RV) stimulation was more effective than translational vertical (TV) stimulation in triggering vertical OKR (Figure 2B, C). In our experiments, binocular rotational stimuli were presented on the left and right screens such that the directions of visual motion were opposite across the two eyes, whereas translational stimuli were presented in the same direction on both screens. Importantly, for both rotational and translational conditions, the spatial and temporal frequencies of the stimuli were identical across eyes.

**Figure 2.**
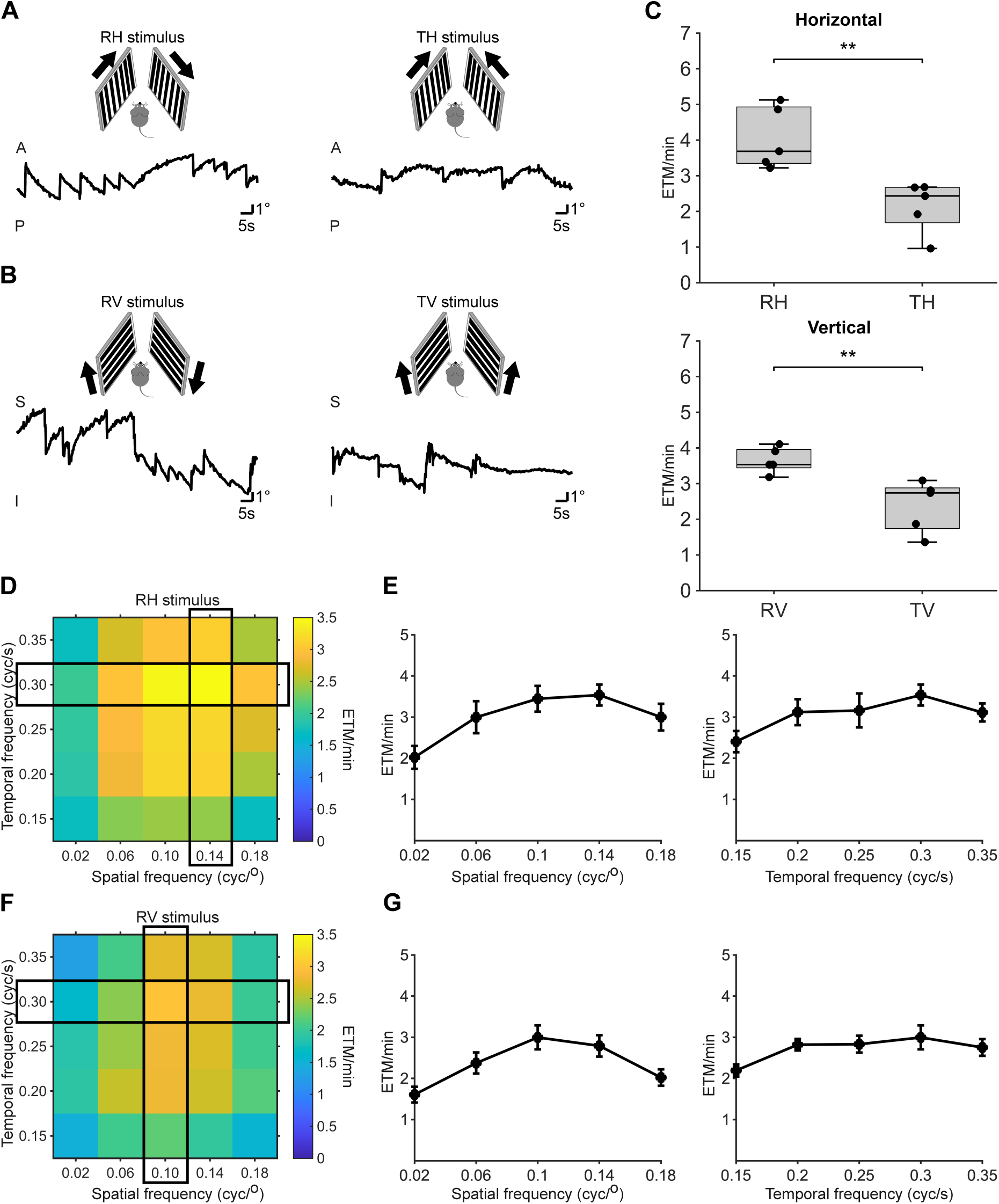
Optimal visual stimulation conditions for eliciting horizontal and vertical OKRs in WT mice. (A) Representative pupil position traces of the left eye in response to horizontal stimulation. Left panel: rotational motion; right panel: translational motion. (B) Representative pupil position traces of the left eye in response to vertical stimulation. Left panel: rotational motion; right panel: translational motion. (C) Quantification of ETMs per minute under different stimulus conditions. Each dot represents one mouse (n = 5 per genotype). (D) Heat map showing average ETMs per minute across temporal–spatial frequency combinations under RH stimulation. The black box marks the optimal temporal and spatial frequency, and the intersection indicates the optimal combination. (E) ETMs curves under RH stimulation. Left: ETMs as a function of spatial frequency at the optimal temporal frequency (0.30 cyc/s, horizontal black box). Right: ETMs as a function of temporal frequency at the optimal spatial frequency (0.14 cyc/°, vertical black box). Data are shown as mean ± SEM (n = 5 mice). (F) Heat map showing average ETMs per minute across temporal–spatial frequency combinations under RV stimulation. The black box marks the optimal temporal and spatial frequency, and the intersection indicates the optimal combination. (G) ETMs curves under RV stimulation. Left: ETMs as a function of spatial frequency at the optimal temporal frequency (0.30 cyc/s, horizontal black box). Right: ETMs as a function of temporal frequency at the optimal spatial frequency (0.10 cyc/°, vertical black box). Data are shown as mean ± SEM (n = 5 mice).

To further explore the optimal temporal and spatial frequencies of visual stimulation that can elicit OKR, we mapped responses across a matrix of TF and SF combinations. Five spatial frequencies (0.02, 0.06, 0.10, 0.14, and 0.18 cycles/degree) were crossed with five temporal frequencies (0.15, 0.20, 0.25, 0.30, and 0.35 cycles/second), generating a total of 25 unique combinations. For each TF × SF combination, we recorded OKR and quantified the efficiency by calculating the average ETMs per minute (Figure 2D, E). For the horizontal OKR, the strongest responses were observed at a temporal frequency of 0.30 cycles/second and a spatial frequency of 0.14 cycles/degree (Figure 2D), with a marked decline in response strength at both higher and lower values (Figure 2E). In contrast, the vertical OKR peaked at a temporal frequency of 0.30 cycles/second and a spatial frequency of 0.10 cycles/degree (Figure 2F), and similarly declined outside this optimal range (Figure 2G). These experiments determined the optimal visual stimulation conditions for horizontal and vertical OKR in mice. In summary, our study identifies a set of visual stimulus parameters that maximize OKR responses and demonstrates that rotational motion is a more effective driver of OKR than translational motion, thereby providing a foundation for future investigations into the neural circuits mediating this reflexive behavior.

### Chrnb2^tm^ mice, like Frmd7^tm^ mice, lack horizontal OKR but retain vertical OKR

Using the optimal stimulus parameters previously determined in WT mice, we next applied the same horizontal and vertical visual stimuli to Chrnb2^tm^ and Frmd7^tm^ mice to assess the impact of these genetic mutations on OKR.

When visual stimuli were presented in the horizontal direction, WT mice exhibited robust horizontal eye tracking movements, as indicated by clear pupil position changes over time (Figure 3A). In striking contrast, both Chrnb2^tm^ and Frmd7^tm^ mice failed to generate any discernible horizontal tracking responses, showing flat pupil position traces regardless of whether the stimulus moved toward the nasal or temporal side (see Video S3 and S5). Under vertical stimulation, all three genotypes produced detectable vertical OKR responses (Figure 3C; see also Video S4 and S6).

**Figure 3.**
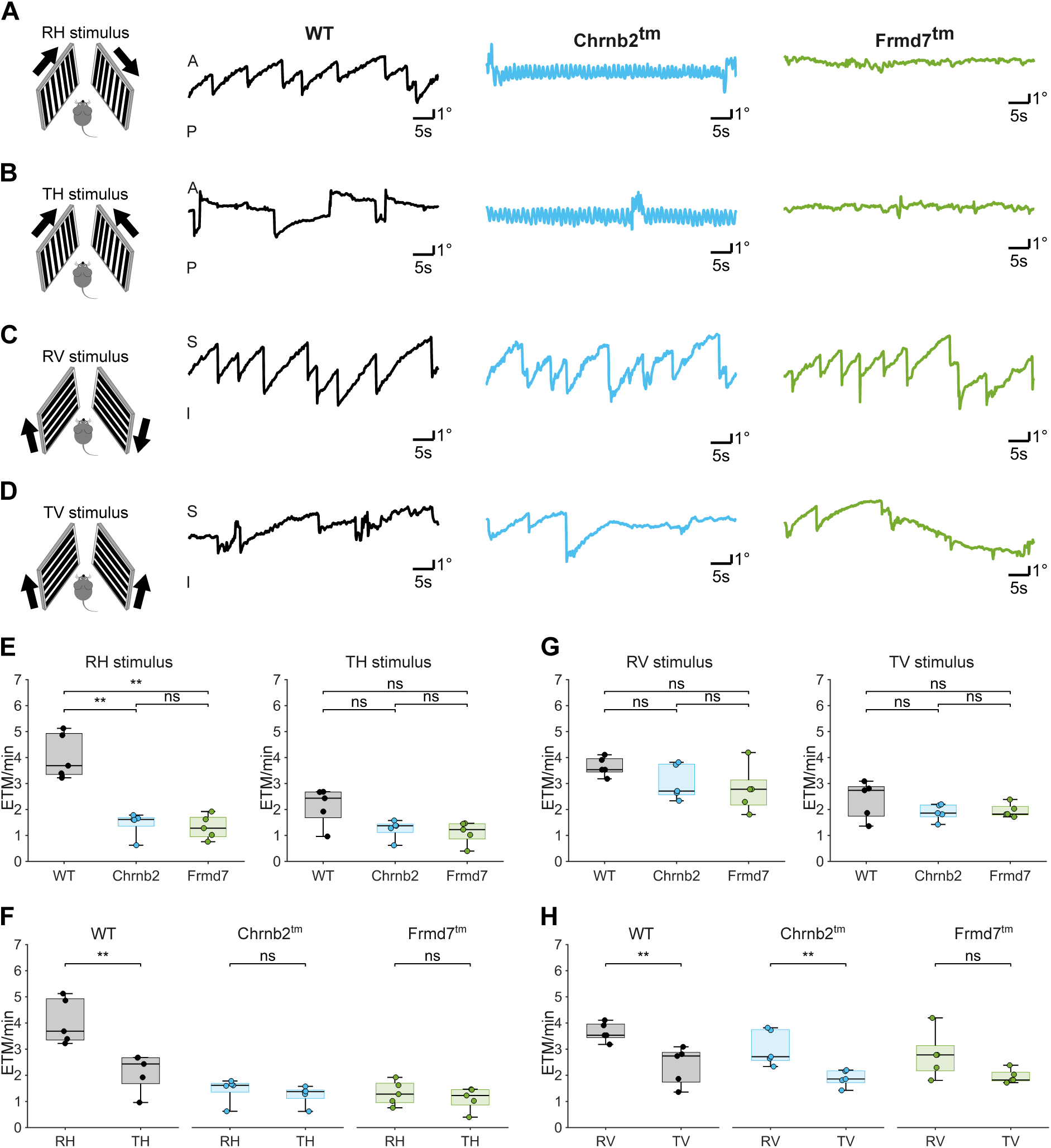
Horizontal OKR is abolished in Chrnb2^tm^ and Frmd7^tm^ mice, whereas vertical OKR remains intact. (A–B) Representative eye movement traces showing pupil position over time in WT, Chrnb2^tm^, and Frmd7^tm^ mice during (A) RH stimulation and (B) TH stimulation. WT mice exhibited robust horizontal OKR under RH and weaker responses under TH, whereas both mutant strains showed no detectable horizontal tracking in either condition. (C–D) Representative vertical OKR traces during (C) RV stimulation and (D) TV stimulation. All genotypes displayed clear vertical responses, with RV stimulation evoking stronger movements than TV. (E–H) Quantification of ETMs per minute under different stimulus conditions. Each dot represents one mouse (n = 5 per genotype). (E) ETMs during RH and TH stimulation. (F) Comparison of ETMs between RH and TH stimulation within each genotype. (G) ETMs during RV and TV stimulation. (H) Comparison of ETMs between RV and TV stimulation within each genotype. Horizontal ETMs were absent in Chrnb2^tm^ and Frmd7^tm^ mice, whereas vertical ETMs were comparable across genotypes.

To quantitatively assess OKR strength, we calculated the number of ETMs per minute during 8 minutes visual stimulation (Figure 3E,G) (See method). In WT mice, horizontal visual stimuli elicited numerous reset saccades, reflecting strong OKR activation. In contrast, both Chrnb2^tm^ and Frmd7^tm^ mice showed a near complete absence of horizontal ETMs, confirming the lack of horizontal OKR (Figure 3E). Notably, vertical ETMs counts were comparable among the three groups, indicating that vertical OKR circuits remain intact despite the loss of horizontal direction selectivity in Chrnb2 ^tm^ mice (Figure 3G). Together, these results demonstrate that Chrnb2^tm^ mice, like Frmd7^tm^ mutants, lack horizontal OKR while retaining vertical OKR.

### Different visual stimulation conditions fail to elicit horizontal OKR in Chrnb2^tm^ mice

To determine whether the absence of horizontal OKR in Chrnb2^tm^ and Frmd7^tm^ mice was specific to a particular set of visual conditions or represented a complete loss of function, we systematically evaluated their eye movement responses under a wide range of visual stimulation paradigms. Specifically, we varied three key stimulus parameters: (1) motion pattern (rotational or translational), (2) frequency (5 spatial frequencies x 5 temporal frequencies), and (3) binocular or monocular stimulation (Figure 5). These responses were directly compared with those of WT mice.

As expected, representative eye movement curves show that in WT mice, RH stimulation evoked strong and sustained horizontal OKR (Figure 3A), whereas TH stimulation induced weaker responses (Figure 3B,F). In contrast, neither Chrnb2^tm^ nor Frmd7^tm^ mice exhibited any horizontal OKR under either motion pattern, indicating a complete loss of horizontal OKR regardless of stimulus pattern (Figure 3A,B,E). When the visual stimuli were presented on the vertical axis, RV stimuli elicited clear vertical OKR in all three genotypes (Figure 3C,G), and TV stimuli also elicited some reflexes, however the responses to TV stimuli were consistently weaker than those evoked by RV stimuli (Figure 3D,H).

Quantification of ETMs across five mice per genotype confirmed these observations. Under horizontal stimulation conditions, WT mice exhibited robust ETMs, whereas no ETMs were detected in Chrnb2^tm^ and Frmd7^tm^ mice. This difference was statistically significant (Figure 4E,F), indicating a loss of horizontal OKR in both mutant lines. In contrast, under vertical stimulation conditions, ETMs were observed in all genotypes with no significant differences across genotypes (Figure 4G,H), suggesting that vertical OKR is largely preserved. Notably, in Frmd7^tm^ mice, there was no significant difference in the intensity of vertical OKR induced by RV and TV stimulation (Figure 3H).

**Figure 4.**
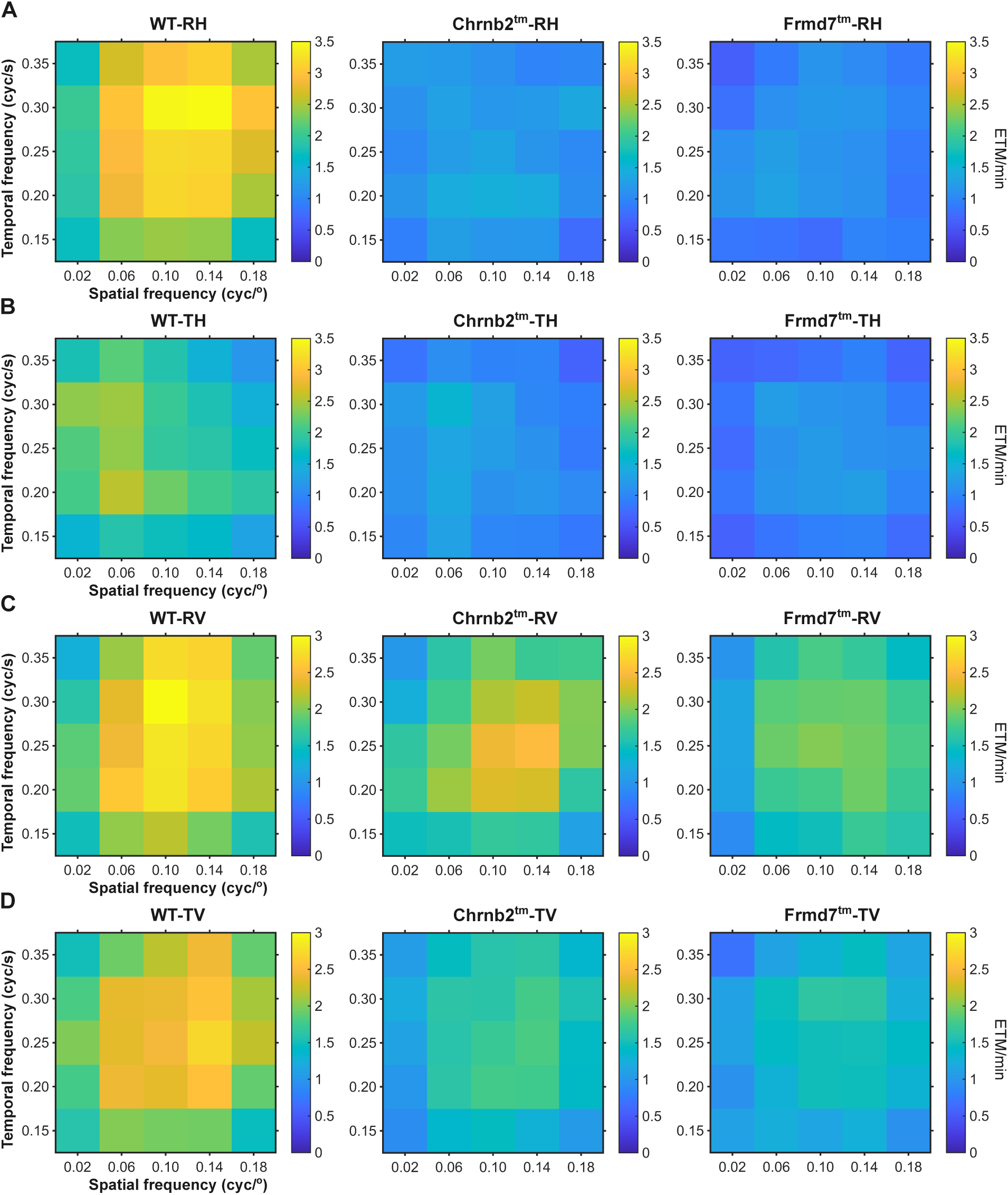
Horizontal OKR is abolished in both Chrnb2^tm^ and Frmd7^tm^ mice, regardless of visual stimulus parameters. (A–B) Heat maps of OKR strength across 25 combinations of spatial frequency (SF) and temporal frequency (TF) during RH stimulation (A) and TH stimulation (B). WT mice displayed maximal horizontal OKR at specific SF-TF combinations under RH, with weaker responses under TH stimulation. No horizontal OKR was exhibited in Chrnb2^tm^ and Frmd7^tm^ mice across all conditions. (C–D) Heat maps of vertical OKR during RV stimulation (C) and TV stimulation (D). All genotypes showed robust vertical responses under RV, with weaker response under TV stimulation (n = 5 per genotype).

We further tested the OKR strength of each movement pattern across 25 spatiotemporal combinations. In WT mice, horizontal OKR was strongest under RH stimulation at specific temporal and spatial frequency combinations (Figure 4A), whereas TH stimulation produced weaker responses across the same parameter matrix (Figure 4B). Consistent with the absence of horizontal OKR, both Chrnb2^tm^ and Frmd7^tm^ mice showed no detectable responses under any frequency combinations in either RH or TH paradigms (Figure 4A,B). In the vertical direction, WT and Chrnb2^tm^ mice showed strong responses to RV stimulation (Figure 4C) but weaker responses to TV stimulation (Figure 4D).

These results demonstrate that Chrnb2^tm^ mice lack horizontal OKR under all the tested stimulus conditions, suggesting a disruption of the neural circuitry responsible for horizontal reflexive eye tracking. Together, these findings are consistent with a role for Chrnb2 in the development of horizontal direction selective retinal circuits.

### Binocular enhancement of vertical OKR is absent in Frmd7^tm^ mice

To further investigate how visual inputs from each eye contribute to OKR, we compared binocular and monocular stimulation conditions in WT, Chrnb2^tm^, and Frmd7^tm^ mice. In binocular trials, both left and right monitors displayed visual motion, whereas in monocular trials, visual motion was presented only to the left side while the opposite monitor displayed a uniform gray field. It is important to note that in this study, “binocular stimulation” refers to presenting visual stimuli on both the left and right screens. However, the stimuli were positioned outside the binocular overlap visual field, meaning that the left eye could not see the stimulus presented on the right screen, and vice versa (see Method).

Under binocular RH stimulation, WT mice exhibited robust and sustained horizontal OKR (Figure 3A). When stimulated by horizontal TH motion, WT mice exhibited weaker responses compared to rotational motion (Figure 3B). When horizontal stimulation was restricted to the left screen, WT mice exhibited reduced OKR compared to binocular RH stimulation, but responses remained comparable to those induced by binocular TH stimulation (Figure 5A,C). This pattern indicates that binocular stimulation under rotational conditions produces enhanced OKR responses compared to monocular stimulation and translational conditions. Horizontal OKR responses were absent across all stimulation conditions in Chrnb2^tm^ mice, suggesting a complete disruption of horizontal OKR circuitry.

**Figure 5.**
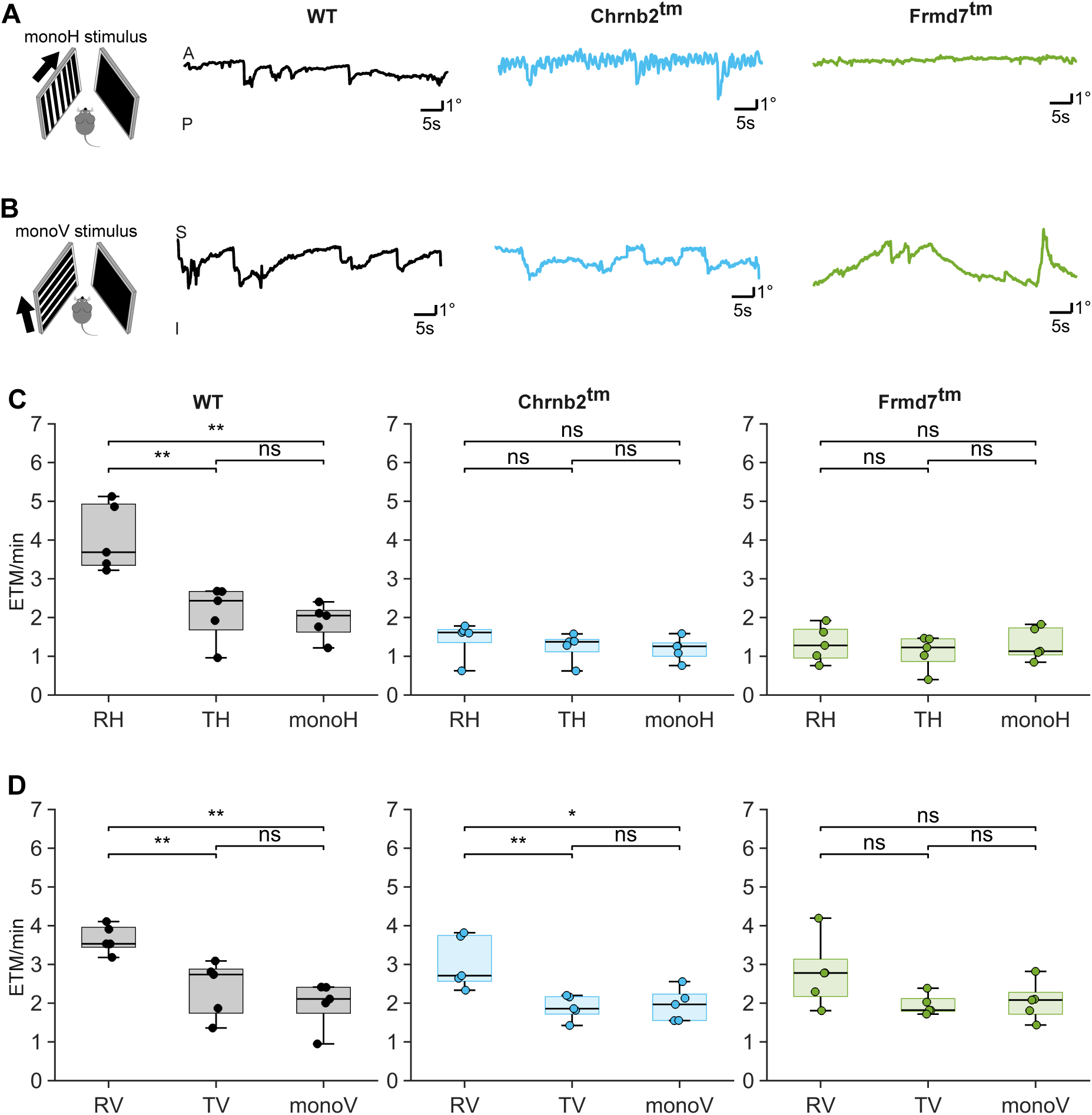
Binocular and monocular stimulation reveal differential properties of horizontal and vertical OKR. (A) Horizontal eye movement traces recorded from the left eye of WT, Chrnb2^tm^, and Frmd7^tm^ mice during monocular stimulation. When the horizontal stimulus was presented only on the left screen, no horizontal OKR was observed in Chrnb2^tm^ and Frmd7^tm^ mice. (B) Vertical eye movement traces during monocular stimulation. (C-D) Quantification of ETMs per minute during monocular stimulation. Each dot represents one mouse (n = 5 per genotype). (C) Horizontal ETMs. WT mice exhibited reduced ETMs under monocular stimulation compared to binocular RH conditions, whereas ETMs remained absent in Chrnb2^tm^ and Frmd7^tm^ mice. (D) Vertical ETMs. WT and Chrnb2^tm^ mice exhibited reduced ETMs under monocular stimulation compared to binocular RV conditions. In Frmd7^tm^ mice, no significant difference was observed between binocular and monocular stimulation. Horizontal OKR remained absent in mutant mice under monocular conditions, whereas vertical OKR was differentially affected across genotypes.

We next applied the same paradigm to vertical motion. During binocular rotational vertical stimulation, all three genotypes exhibited clear vertical OKR (Figure 3C), confirming that vertical pathways remain intact in both mutant lines. In contrast, responses under translational vertical stimulation were weaker (Figure 3C). These data indicate that rotational stimuli are generally more effective than translational stimuli for eliciting vertical OKR, consistent with our findings for horizontal motion. When vertical stimulation was restricted to the left screen, weak vertical OKR was elicited in all three genotypes (Figure 5B).

To summarize these effects, we quantified the mean ETMs per minute of each mouse. In WT mice, ETMs counts were significantly higher under binocular RH stimulation compared to both TH and monocular horizontal (monoH) conditions (Figure 5C). A similar pattern was observed for vertical stimulation; binocular RV stimulation elicited higher ETMs counts than TV and monocular vertical (monoV) conditions (Figure 5D). WT mice show enhanced OKR responses specifically under binocular rotational stimulation. Although ETMs counts during binocular vertical stimulation were comparable between mutant and WT mice (Figure 3G), their response patterns differed markedly under different stimulation conditions. Chrnb2^tm^ mice exhibited comparable response patterns to WT mice, with significantly higher ETMs counts under RV stimulation and no difference between TV and monoV conditions. In contrast, Frmd7^tm^ mice showed no significant differences across the three stimulation conditions, with comparable ETMs counts under RV, TV, and monoV stimulation (Figure 5D). These findings imply that, although both mutant strains retain vertical OKR, binocular enhancement under rotational conditions is preserved in Chrnb2^tm^ mice but absent in Frmd7^tm^ mice.

### Chrnb2^tm^ mice exhibit abnormal horizontal oscillatory eye movements

To examine horizontal eye movement control, we analyzed pupil position dynamics in WT, Chrnb2^tm^, and Frmd7^tm^ mice under identical recording conditions without visual stimulation. While WT and Frmd7^tm^ mice showed relatively stable horizontal eye positions, Chrnb2^tm^ mice exhibited pronounced oscillatory eye movements confined to the horizontal axis (Figure 6A). Spatial density maps of pupil position revealed a broadened distribution in Chrnb2^tm^ mice compared with the compact distributions observed in WT and Frmd7^tm^ mice (Figure 6B), indicating increased instability. Consistent with the broadened pupil position distributions observed in the spatial density maps, quantitative analysis revealed a significant increase in eye drift in Chrnb2^tm^ mice compared with WT and Frmd7^tm^ controls (Figure 6C). Fast Fourier Transform (FFT) analysis further revealed that horizontal eye movements in Chrnb2^tm^ mice were dominated by rhythmic oscillations at approximately 10 Hz (Figure 6D). This oscillatory component was persistent over time and was not observed in WT or Frmd7^tm^ mice. Consistently, power spectral analysis demonstrated an elevated power at approximately 10 Hz in Chrnb2^tm^ mice, whereas WT and Frmd7^tm^ mice exhibited spectra dominated by low frequency components without a distinct peak (Figure 6D). Correspondingly, quantitative analysis revealed that chrnb2^tm^ mice exhibited significantly stronger power at 10 Hz. (Figure 6F). We further analyzed vertical eye movements in the absence of visual stimulation. In contrast to the horizontal axis, no distinct frequency bands or power peaks were observed in the vertical direction across the three mouse strains (Figure 6E). In Chrnb2^tm^ mice, the power at 10 Hz was significantly stronger during horizontal eye movements than during vertical eye movements (Figure 6G). These findings indicate that Chrnb2^tm^ mice exhibit ∼10 Hz oscillatory eye movements that are predominantly along the horizontal axis.

**Figure 6.**
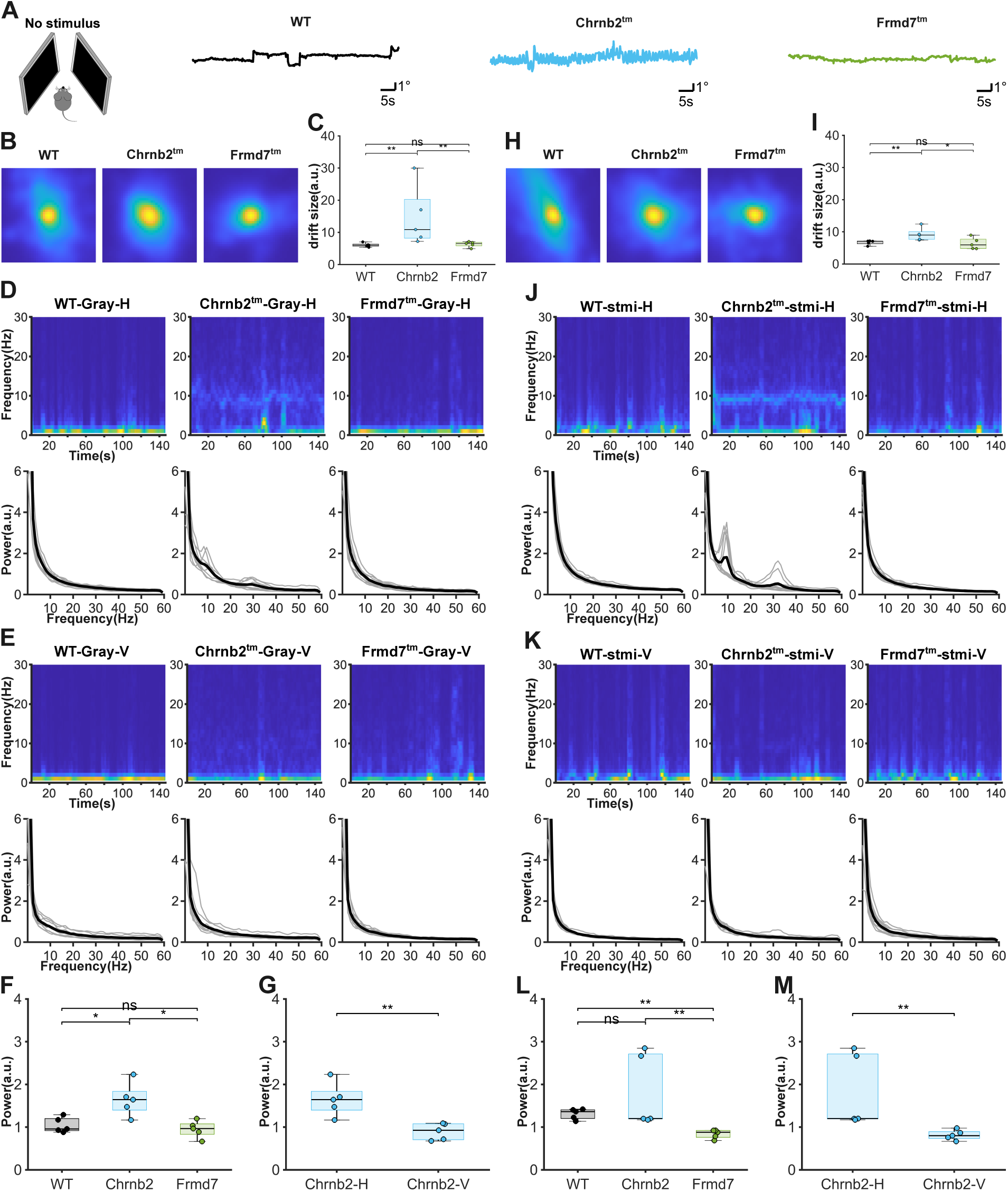
Chrnb2^tm^ mice exhibit abnormal horizontal oscillatory eye movements. (A) Changes in pupil position over time under no stimulated conditions (gray) in WT, Chrnb2^tm^, and Frmd7^tm^ mice. (B) Spatial density maps of horizontal pupil position in WT, Chrnb2^tm^, and Frmd7^tm^ mice under no stimulated conditions. WT and Frmd7^tm^ mice show compact distributions centered around the mean eye position, whereas Chrnb2^tm^ mice display a broadened, indicating decrease eye instability. (C) Quantification of eye drift. Each dot represents one mouse (n = 5 per genotype). Chrnb2^tm^ mice exhibited significantly increased eye drift compared with WT and Frmd7^tm^ mice, indicating elevated eye position instability. (D) Top: Representative figure from Fast Fourier Transform (FFT) analysis of horizontal eye movements. In the absence of visual stimulation, Chrnb2^tm^ mice exhibit prominent rhythmic activity at approximately 10 Hz that persists over time. Bottom: Power spectral of eye movements signals obtained by FFT. The gray curve represents a single record, with 2-3 records per animal. The black curve represents the average of multiple records. Chrnb2^tm^ mice display elevated power at approximately 10 Hz frequency compared with WT and Frmd7^tm^ mice, which lack a distinct spectral peak and are dominated by low-frequency components. (E) Same analysis as in (D) but for vertical eye movements. (F) Quantification of eye drift power at 10 Hz. Each dot represents one mouse (n = 5 per genotype). Chrnb2^tm^ mice exhibited significantly higher values compared with WT and Frmd7^tm^ mice (*p < 0.05), whereas WT and Frmd7^tm^ mice did not differ significantly. (G) Comparison of power at 10 Hz during horizontal (Chrnb2-H) and vertical (Chrnb2-V) eye movement in Chrnb2^tm^ mice. Horizontal eye movement exist significantly larger power (p < 0.01). (H) Spatial density maps of horizontal pupil position in WT, Chrnb2^tm^ mice, and Frmd7^tm^ mice during horizontal visual stimulation. (I) Quantification of eye drift under horizontal visual stimulation. Chrnb2^tm^ mice exhibited significantly increased eye drift compared with WT and Frmd7^tm^ mice. (J) Top: Representative figure from FFT analysis of horizontal eye movements during horizontal visual stimulation. Chrnb2^tm^ mice exhibit prominent rhythmic activity at approximately 10 Hz that persists over time. Bottom: Power spectra of eye movement signals obtained by FFT. Chrnb2^tm^ mice display elevated power around 10 Hz compared with WT and Frmd7^tm^ mice. (K) Same analysis as in (J) but for vertical eye movements. (L) Quantification of eye movement power at 10 Hz under horizontal visual stimulation. (M) Comparison of power at 10 Hz during horizontal and vertical eye movements in Chrnb2^tm^ mice under visual stimulation.

To determine whether these oscillatory eye movements persist under visual stimulation, we performed the same analyses during horizontal visual stimulation. Spatial density maps of pupil position revealed a broadened distribution in Chrnb2^tm^ mice compared with the compact distributions observed in WT and Frmd7^tm^ mice (Figure 6H). Quantitative analysis confirmed this (Figure 6I), indicating Chrnb2^tm^ mice also exhibited increased horizontal instability under visual stimulation, consistent with results obtained under unstimulated conditions. FFT analysis further revealed that horizontal eye movements in Chrnb2^tm^ mice were dominated by rhythmic oscillations at approximately 10 Hz (Figure 6J), similar to the oscillatory pattern observed in the absence of visual stimulation. Consistently, power spectral analysis demonstrated an elevated power at approximately 10 Hz in Chrnb2^tm^ mice, whereas WT and Frmd7^tm^ mice exhibited spectra dominated by low-frequency components without a distinct peak (Figure 6J). Quantitative analysis revealed that the power at 10 Hz was significantly higher in Chrnb2^tm^ mice than in Frmd7^tm^ mice, while no significant difference was detected between Chrnb2^tm^ and WT mice. WT mice also showed significantly higher 10 Hz power compared with Frmd7^tm^ mice (Figure 6L). This may reflect stimulus-related rhythmic components in WT eye movements under horizontal visual stimulation. We further analyzed vertical eye movements during horizontal visual stimulation. No distinct frequency bands or power peaks were observed in the vertical direction across the three mouse strains (Figure 6K). In Chrnb2^tm^ mice, the power at 10 Hz remained significantly stronger during horizontal eye movements than during vertical eye movements (Figure 6M). In summary, these results indicate that Chrnb2^tm^ mice exhibit oscillatory eye movements, which persist under visual stimulation, and are primarily limited to the horizontal direction. Together, these results indicate that loss of Chrnb2 selectively disrupts horizontal eye movement stability, leading to high-frequency oscillatory movements.

## Discussion

In this study, we systematically compared the horizontal and vertical eye movements of WT, Chrnb2^tm^, and Frmd7^tm^ mice using a customized behavioral recording system capable of tracking horizontal and vertical pupil movements (Figures 1-6). Our results demonstrate that both Chrnb2^tm^ and Frmd7^tm^ mice completely lack horizontal OKR while maintaining vertical OKR across a broad range of stimulus conditions. These findings establish Chrnb2 as an essential gene for the development of horizontal direction selective retinal circuits and their downstream pathways, paralleling the role of Frmd7. Consistent with previous reports(Bistrong et al., 2025), Chrnb2^tm^ mice lacked horizontal OKR while preserving vertical responses. Importantly, we found they exhibited spontaneous horizontal oscillatory eye movements, which were absent in Frmd7^tm^ mice. These findings suggest that the horizontal deficit in Chrnb2^tm^ mice is accompanied by circuit instability, contrasting with the pure loss of directional tuning inherent in the Frmd7 phenotype. In addition, WT mice showed enhanced OKR responses under binocular rotational stimulation in both horizontal and vertical axes. This enhancement was preserved in Chrnb2^tm^ mice along the vertical axis, but was absent in Frmd7^tm^ mice along the same axis, indicating a selective loss of binocular enhancement in the Frmd7^tm^ mice.

### Chrnb2 and retinal DS circuits

Our findings highlight the critical role of β2-nAChRs in shaping direction-selective circuits in the retina. Previous work has shown that β2-nAChRs, encoded by Chrnb2, are required for cholinergic retinal waves that refine visual circuits during development(Bansal et al., 2000; C. Sun et al., 2008). Disruption of these receptors results in abnormal circuit maturation and selective impairment of DSGCs, with horizontal preferring DSGCs more severely affected than vertical preferring subtypes(Bistrong et al., 2025). This difference may reflect evolutionary specialization; horizontal image stabilization is essential for locomotion and navigation in rodents, whereas vertical stabilization may rely on partially inputs from other visual pathways, such as the superior colliculus or vestibular inputs, allowing vertical OKR to persist despite partial retinal dysfunction(Simpson, 1984; Yonehara et al., 2011).

The complete absence of horizontal OKR in Chrnb2^tm^ mice therefore reflects the failure of horizontal DS circuits to provide appropriate input to the AOS, which normally drives horizontal compensatory eye movements via the NOT and DTN(Dhande et al., 2013; Simpson, 1984). An alternative possibility is that the absence of horizontal OKR arises from differences in stimulus conditions rather than an underlying circuit deficit. To address this, we systematically explored a broad range of visual stimulus parameters, including spatial frequence (0.02–0.18 cycles/degree) and temporal frequencies (0.15–0.35 cycles/s). In addition, we tested multiple stimulus configurations, including binocular rotational and translational motion as well as monocular stimulation. However, Chrnb2^tm^ mice failed to exhibit any detectable horizontal OKR under any condition tested (Figures 3,4), arguing against the possibility that the deficit arises from inappropriate stimulus tuning or stimulus configuration. This strict dependency highlights the NOT is a key for horizontal image stabilization and positions horizontal DSGCs as a critical retinal circuit for this behavior.

### A brainstem model for binocular integration of rotational optic flow and its selective disruption in Frmd7^tm^ mice

In our paradigm, rotational stimulation was more effective than translational stimulation in driving robust OKR (Figure 2), suggesting that the visuomotor system is particularly sensitive to motion patterns characteristic of self-rotation. This distinction aligns with the fundamental organization of optic flow processing; opposite direction motion across the two visual hemifields defines rotational optic flow, a primary cue for detecting head rotation, whereas same direction motion defines translational optic flow, which typically reflects forward or lateral body displacement.

Behavioral and physiological studies across species consistently show that rotational optic flow elicits stronger and higher gain optokinetic responses than translational optic flow, reflecting the greater demands of stabilizing gaze during head rotation(Cullen, 2012; Tian et al., 2007; Wang et al., 2019; Nalbach et al., 1993). For example, mammalian studies demonstrate that horizontal rotational OKN exhibits higher gain and distinct kinematics compared with translational OKN(Tian et al., 2007), and zebrafish pretectal and tectal circuits contain separate neural populations selectively tuned to rotational versus translational flow fields, with rotational responses generally stronger and more informative for stabilizing self-motion(Wang et al., 2019). Similarly, classic split field studies in crabs show that binocular motion in opposite directions strongly enhances optokinetic gain, whereas same direction binocular motion leads to weaker or more saturating responses(Nalbach et al., 1993). Together, these results indicate that the rotation and translation asymmetry in our data represents a conserved, cross species principle of optic flow processing. Subcortical structures homologous to the mammalian AOS, such as the avian nBOR, further demonstrate selective tuning to rotational and translational optic flow(1999), highlighting that global motion patterns are segregated early in the visuomotor hierarchy. In mice, binocular integration of retinal motion contributes to the formation of rotation and translation selective optic flow representations in the visual cortex, and these representations have been reported to be reduced in Frmd7^tm^ mice(Rasmussen et al., 2021). These findings complement our behavioral evidence that rotational stimulation provides a more potent drive for OKR than translational stimulation.

Although AOS nuclei are largely monocular, the visual motion signals they convey ultimately enter brainstem pathways that operate on binocular optic flow, because global visual motion during natural behavior is experienced simultaneously through both eyes. Convergent evidence shows that the vestibular nuclear complex, particularly the medial vestibular nucleus, serves as a major hub where retinal slip signals are combined with bilateral vestibular inputs to estimate self-motion and generate premotor commands for gaze stabilization. In this context, rotational optic flow may naturally engages the bilateral push–pull architecture of the vestibular nuclei, potentially producing a stronger premotor drive than translational optic flow, whose binocular input is symmetric and largely redundant(Cullen, 2012; Medrea & Cullen, 2013; Wibble et al., 2022). Although direct experimental evidence for rotation-specific binocular interactions in the vestibular nuclei is still lacking, our results are consistent with the possibility that such mechanisms may contribute to the enhanced OKR observed under rotational stimulation. Future recordings from vestibular nuclei under opposite and same direction binocular stimulation will be necessary to test this hypothesis.

This framework also offers insight into the selective loss of vertical binocular enhancement in Frmd7^tm^ mice (Figure 5). Developmental expression studies show that FRMD7 is expressed not only in retinal direction-selective circuits but also in tissues associated with VOR/OKR pathway formation, including the vestibular apparatus, vestibulocochlear ganglion, hindbrain ventricular zones, brainstem, and cerebellum(Betts-Henderson et al., 2010; Tarpey et al., 2006; Thomas et al., 2014; Yonehara et al., 2016). Combined with evidence that FRMD7 regulates CASK-dependent actin remodeling and neurite outgrowth in oculomotor pathways(2010; Watkins et al., 2013), these findings raise the possibility that Frmd7 deficiency may affect retinal direction selectivity and the maturation or weighting of binocular-vestibular convergence in brainstem premotor circuits. Such a disruption would selectively reduce binocular enhancement during rotational optic flow, while leaving the weaker translational responses relatively unaffected. In contrast, Chrnb2 primarily influences retinal-wave dependent refinement of retinal projections, with no compelling evidence for its role in brainstem visual-vestibular circuit development at the moment(Bansal et al., 2000; Grubb & Thompson, 2004; McLaughlin et al., 2003), which likely explains the preserved vertical binocular enhancement in Chrnb2^tm^ mice.

### Oscillatory pupil movements in Chrnb2^tm^ mice

Unexpected phenotype observed in Chrnb2^tm^ mice was spontaneous pupil oscillations (Figure 6). These oscillations occurred both during visual stimulation and in the absence of visual stimulation. One plausible explanation is that disruption of cholinergic signaling in the developing retina, resulting from the loss of β2-nAChRs, leads to abnormal synchronization or rhythmic activity within retinal circuits that is subsequently transmitted to downstream eye movement centers. Starburst amacrine cell mediated cholinergic retinal waves are known to play a critical role in shaping the maturation and patterning of DSGC circuits(Bansal et al., 2000; Elstrott et al., 2008), and alterations in wave dynamics can generate atypical correlated activity in retinal outputs. If ON-DSGCs become abnormally synchronized or intrinsically rhythmic as a consequence of disrupted early activity, such oscillatory retinal output could be relayed via the accessory optic system and pretectal pathways to drive rhythmic pupil or eye movements(Borowska et al., 2011; Trenholm et al., 2011).

Recent studies further indicate that the development of retinal direction-selective circuits depends critically on early spontaneous activity and proceeds in an axis-specific manner(Bistrong et al., 2025). Specifically, β2-nAChRs encoded by Chrnb2 are required for the proper maturation of horizontal, but not vertical, direction selectivity in ON-DSGCs(Huberman et al., 2008). In Chrnb2^tm^ mice, the asymmetric inhibitory inputs that normally confer horizontal direction tuning are weakened or spatially misaligned, indicating impaired activity-dependent refinement rather than a complete loss of direction-selective cell identity(Bistrong et al., 2025).

Together, these findings suggest that loss of Chrnb2 does not simply eliminate the directional signal in ON-DSGCs, but instead alters its output pattern, potentially making it contain irregular or rhythmic components. Such aberrant directional signaling could destabilize visuomotor feedback loops and contribute to the abnormal pupil or eye oscillations observed in these animals. This interpretation is consistent with previous work showing that retinal direction-selective signals are a major driver of reflexive eye movements and that perturbations of retinal activity patterns can profoundly influence oculomotor behavior(Oyster & Barlow, 1967; Wei & Feller, 2011).

By contrast, although Frmd7^tm^ mice lack horizontal direction selectivity and exhibit a loss of horizontal OKR, they did not display such pupil oscillations in our recordings(Yonehara et al., 2016). This difference suggests that loss of horizontal direction tuning alone is insufficient to generate the oscillatory phenotype. Instead, oscillations may require abnormal retinal activity patterns, such as disrupted developmental retinal waves, which are unique to the consequences of Chrnb2 loss. The difference is also relevant to human disease: FRMD7 mutations cause congenital nystagmus in humans(Huang et al., 2022; Tarpey et al., 2006), yet the pathway from retinal circuit abnormality to oscillatory oculomotor output likely involves additional factors, such as species-specific differences in central circuit organization, plasticity or compensatory mechanisms(Dell’Osso & Jacobs, 2002; Thurtell & Leigh, 2011).

### Rotational stimulation more effective than translational stimulation

Rotational stimulation robustly elicited both horizontal and vertical OKRs, whereas translational stimulation induced only weak responses. The continued advantage of this motion simulating yaw and roll axis rotations indicates the AOS is preferentially tuned to rotational optic flow, likely reflecting its role in stabilizing vision during natural behaviors involving head rotation. In contrast, during forward movement, translational inputs are less effective at driving OKR, possibly as a mechanism to avoid excessive reflexive eye movements. Previous studies have emphasized rotation as the classical stimulus for horizontal OKR (Collewijn, 1969), and our results extend this conception by demonstrating that rotational dominance applies not only to horizontal reflexes mediated by the NOT/DTN, but also to vertical reflexes mediated by the MTN.

Taken together, our findings reveal that Chrnb2 is crucial for the development of horizontal DS circuits and their behavioral output in the form of horizontal OKR, paralleling the role of Frmd7. Furthermore, WT mice exhibited enhanced OKR responses under binocular rotational stimulation, an effect that was preserved in Chrnb2^tm^ mice but absent in Frmd7^tm^ mice, indicating a selective loss of binocular enhancement in the Frmd7^tm^ mice. In addition, the discovery of spontaneous oscillatory pupil movements in Chrnb2^tm^ mice identifies a previously unrecognized phenotype that provides new insight into the diversity of circuit level mechanisms leading to abnormal oculomotor behaviors. By directly linking specific genetic deletions to both circuit deficits and behavioral outcomes, our work contributes to a mechanistic understanding of how retinal circuits support visual stability and how their disruption can result in distinct pathological phenotypes.

### Limitations of the study

While our study provides a detailed behavioral characterization of horizontal and vertical OKR in WT, Chrnb2^tm^, and Frmd7^tm^ mice, it is limited by the lack of direct electrophysiological recordings from retinal or brainstem circuits. Consequently, the cellular and synaptic mechanisms underlying the observed phenotypes remain to be confirmed. Future studies combining circuit-level recordings with varied visual stimulation will be important to fully elucidate the mechanistic basis of these OKR phenotypes.

## Resource availability

Lead contact: Further information and requests for resources, reagents, and data should be directed to and will be fulfilled by the lead contact, [Keisuke Yonehara].

Materials availability: Wild-type (C57BL/6J) mice, Chrnb2^tm^ mice (Chrnb2^tm1a(EUCOMM)Hmgu^, EMMA ID: EM:07867), and Frmd7^tm^ mice (Frmd7^tm1a(KOMP)Wtsi^, MGI:4455520) used in this study were obtained from public repositories. All strains are available from the respective repositories upon reasonable request.

Data and code availability: All raw behavioral OKR data and MATLAB analysis scripts generated during this study are available from the lead contact upon request.

## Acknowledgements

We thank Simon Arvin for developing the Eyeloop software for pupil tracking and for guidance on its implementation. We thank Monica Dahlstrup Sietam for helpful discussions and advice during the development of the OKR tracking platform. We also thank all members of the Yonehara laboratory for valuable discussions and support.

This work was supported by grants from KAKENHI (20K23377; 22K21353; 23H04241; 24H02311), Chugai Foundation for Innovative Drug Discovery Science, Daiichi Sankyo Foundation of Life Science, Mochida Memorial Foundation for Medical and Pharmaceutical Research, Mitsubishi Foundation, Toray Science Foundation, and Naito Foundation to K.Y.

## Author contributions

Conceptualization, K.Y.; Funding acquisition, K.Y.; Investigation, J.Q.; Data curation, J.Q.; Formal analysis, A.M.; Writing – original draft, J.Q.; Writing – review & editing, K.Y. and A.M.; Supervision, K.Y.

## Methods

### Experimental animals

For wild-type mice (C57BL/6J), 3 males and 2 females were used. Chrnb2^tm1a(EUCOMM)Hmgu^ mice (EMMA ID: EM:07867) were obtained from the European Mouse Mutant Archive (EMMA, Infrafrontier) and crossed with female Sox2-Cre delete mice to excise the loxP-flanked exon 5 of the Chrnb2 gene. The resulting germline deletion allele is referred to as Chrnb2^tm^. For Chrnb2^tm^ mice, homozygous 3 males and 2 females were used. Frmd7^tm1a(KOMP)Wtsi^ mice (MGI:4455520) were obtained from the Knockout Mouse Project (KOMP) Repository and crossed with female Sox2-Cre mice to excise the loxP-flanked exon4, generating Frmd7^tm^ mice. For Frmd7^tm^ mice, 2 homozygous females and 3 hemizygous males were used. Mice were maintained in a 12-hour light/12-hour dark cycle environment with free access to food and water. All animal experiments were performed according to standard ethical guidelines and were approved by the National Institute of Genetics (R7-1).

### Head-fixation surgery

Mice were anaesthetized by intraperitoneal injection of a mixture of butorphanol (5 mg/kg), midazolam (4 mg/kg), and medetomidine (0.5 mg/kg). Adequate anesthesia was confirmed by absence of the pedal withdrawal reflex. Throughout the surgical procedure, mice were maintained on a temperature-controlled heating pad, and 0.3% ofloxacin eye ointment (Santen Pharmaceutical) was applied to the eyes to prevent dehydration and corneal damage. After the scalp was incised along the midline to expose the skull, the skull surface was gently roughened with a scalpel blade to improve adhesion. Screws were then fixed to the skull using cyanoacrylate adhesive (Super Glue Precision, Loctite) and reinforced with dental cement (Sun Medical). At the end of surgery, mice received atipamezole (0.75 mg/kg, i.p.) to reverse anesthesia and were allowed to recover in a heated cage until fully awake. Surgeries were completed at least three days before experiments.

### Experimental setup for OKR tests

Head-fixed mice were placed in an open restraining tube positioned on a platform. Two monitors (52 × 30 cm) were placed parallel to the animal’s midline at a distance of 15 cm from the eyes, with each monitor corresponding to one eye. For each eye, visual stimuli spanned an azimuth range of 30°–150° and an elevation range of −45° to +45°, centered on the optic axis of that eye. A 45° hot mirror was positioned on the left side of the mouse to allow simultaneous visual stimulation and eye tracking(Yonehara et al., 2009). A near-infrared light source (SLS-0208-B LED Spot Light) was placed posterior-right to the animal to illuminate the eye. A CCD camera (Allied Vision Guppy Pro F-031, 1/4″ monochrome sensor) was positioned on the left side of the mouse and connected to a PC via a PCIe frame-grabber (ADLINK FIW62). Video frames were streamed in real time at ∼120 Hz into EyeLoop(Arvin et al., 2021) using the Vimba-based importer (vimba.py). All records in the experiment were taken from the left eye of the mouse.

### Visual stimulation

Visual stimuli were generated using Psychtoolbox(Brainard, 1997) on MATLAB (MathWorks) and presented on screens positioned on both sides of the animal. To evoke the horizontal OKR in WT and mutant mice, drifting gratings simulating yaw-axis rotation and forward–backward translation were presented. Gratings drifted at 0° and 180° (0.14 cycles/°, 0.30 cycles/s). To evoke the vertical OKR, drifting gratings simulating roll-axis rotation and up–down translation were presented. Gratings drifted at 90° and 270° (0.10 cycles/°, 0.25 cycles/s). Drifting direction alternated every 2 min (total 8 min). For monocular stimulation, gratings were presented only on the left screen. Each mouse was recorded 2–3 times. For experiments using 25 frequency combinations, direction alternated every 1 min (total 4 min), and stimuli were presented binocularly.

### Statistical analysis

Statistical analyses were performed using MATLAB. Data are presented as box-and-whisker plots, where the center line indicates the median, and the box represents the interquartile range. Whiskers extend to the minimum and maximum values. Dots represent the mean value for an individual animal, calculated from 2–3 repeated recordings. As data distributions were not assumed to be normal, non-parametric statistical tests were used. Comparisons between two independent groups were performed using two-tailed Mann–Whitney U tests. For multiple group comparisons, pairwise Mann–Whitney U tests were conducted between groups as indicated. Statistical significance was defined as p < 0.05. Significance levels are denoted as p < 0.05 (*), p < 0.01 (**), and p < 0.001 (***); non-significant differences are labeled as “ns”.

### Quantification of the optokinetic reflex

The OKR were quantified as the number of eye tracking movements (ETMs)(Cahill & Nathans, 2008). To detect ETMs, first we computed the derivative of positions of the pupil (*Δt_r_*), and a threshold:

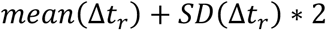

was set for each data. Next, the detected event candidates above threshold were screened based on the template matching. The templates were established by using the datasets of representative OKR of WT mice during visual stimulation. We computed cross-correlation (*C_r_*) between individual event candidates and the templates:

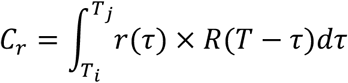

The events with *C_r_* above 0.8 were determined as ETMs.

### Quantification of oscillatory eye movements

To quantify the stability of pupil, we first mapped temporal changes of the pupil locations on 2D spaces with a smoothing by moving-average. Next, the maps were smoothed by 2D Gaussian filters and centered by a peak location to obtained an average spatial maps for WT and mutant population. To quantify the oscillation in the pupil locations, we used Fast Fourier Transform (FFT) and FFT moving average with 50 seconds time window. The oscillatory power was determined as scores at 10 and 30 Hz.

## Supplemental information

Video S1. Horizontal OKR in WT mice (Related to Figure 1B,C). Representative recording of a head-fixed WT mouse under RH stimulation. All recordings were obtained from the left eye.

Video S2. Vertical OKR in WT mice (Related to Figure 1B,D). Representative recording of a head-fixed WT mouse under RV stimulation.

Video S3. Horizontal eye movement in Chrnb2^tm^ mice (Related to Figure 3A). Representative recording of a head-fixed Chrnb2^tm^ mouse under RH stimulation.

Video S4. Vertical OKR in Chrnb2^tm^ mice (Related to Figure 3C). Representative recording of a head-fixed Chrnb2^tm^ mouse under RV stimulation.

Video S5. Horizontal eye movement in Frmd7^tm^ mice (Related to Figure 3A). Representative recording of a head-fixed Frmd7^tm^ mouse under RH stimulation.

Video S6. Vertical OKR in Frmd7^tm^ mice (Related to Figure 3C). Representative recording of a head-fixed Frmd7^tm^ mouse under RV stimulation.

## Reference

Ackman, J. B., Burbridge, T. J., & Crair, M. C. (2012). Retinal waves coordinate patterned activity throughout the developing visual system. Nature, 490(7419), 219–225. 10.1038/nature11529

Arvin, S., Rasmussen, R. N., & Yonehara, K. (2021). EyeLoop: An Open-Source System for High-Speed, Closed-Loop Eye-Tracking. Frontiers in Cellular Neuroscience, 15, 779628. 10.3389/fncel.2021.779628

Balaban, C. D. (1988). Distribution of inferior olivary projections to the vestibular nuclei of albino rabbits. Neuroscience, 24(1), 119–134. 10.1016/0306-4522(88)90317-x

Bansal, A., Singer, J. H., Hwang, B. J., Xu, W., Beaudet, A., & Feller, M. B. (2000). Mice lacking specific nicotinic acetylcholine receptor subunits exhibit dramatically altered spontaneous activity patterns and reveal a limited role for retinal waves in forming ON and OFF circuits in the inner retina. The Journal of Neuroscience: The Official Journal of the Society for Neuroscience, 20(20), 7672–7681. 10.1523/JNEUROSCI.20-20-07672.2000

Barlow, H. B., & Hill, R. M. (1963). Selective Sensitivity to Direction of Movement in Ganglion Cells of the Rabbit Retina. Science, 139(3553), 412–414. 10.1126/science.139.3553.412

Betts-Henderson, J., Bartesaghi, S., Crosier, M., Lindsay, S., Chen, H.-L., Salomoni, P., Gottlob, I., & Nicotera, P. (2010). The nystagmus-associated FRMD7 gene regulates neuronal outgrowth and development. Human Molecular Genetics, 19(2), 342–351. 10.1093/hmg/ddp500

Bistrong, K., Somaiya, R. D., Liang, E. Y., Smith, B. E., & Feller, M. B. (2025). Activity-dependent development of synaptic circuits mediates direction selectivity in an axis-specific manner. Cell Reports, 44(7). 10.1016/j.celrep.2025.115897

Borowska, J., Trenholm, S., & Awatramani, G. B. (2011). An intrinsic neural oscillator in the degenerating mouse retina. The Journal of Neuroscience: The Official Journal of the Society for Neuroscience, 31(13), 5000–5012. 10.1523/JNEUROSCI.5800-10.2011

Brainard, D. H. (1997). The Psychophysics Toolbox. Spatial Vision, 10(4), 433–436.

Burbridge, T. J., Xu, H.-P., Ackman, J. B., Ge, X., Zhang, Y., Ye, M.-J., Zhou, Z. J., Xu, J., Contractor, A., & Crair, M. C. (2014). Visual circuit development requires patterned activity mediated by retinal acetylcholine receptors. Neuron, 84(5), 1049–1064. 10.1016/j.neuron.2014.10.051

Cahill, H., & Nathans, J. (2008). The optokinetic reflex as a tool for quantitative analyses of nervous system function in mice: Application to genetic and drug-induced variation. PloS One, 3(4), e2055. 10.1371/journal.pone.0002055

Collewijn, H. (1969). Optokinetic eye movements in the rabbit: Input-output relations. Vision Research, 9(1), 117–132. 10.1016/0042-6989(69)90035-2

Cullen, K. E. (2012). The vestibular system: Multimodal integration and encoding of self-motion for motor control. Trends in Neurosciences, 35(3), 185–196. 10.1016/j.tins.2011.12.001

Dell’Osso, L. F., & Jacobs, J. B. (2002). An expanded nystagmus acuity function: Intra- and intersubject prediction of best-corrected visual acuity. Documenta Ophthalmologica. Advances in Ophthalmology, 104(3), 249–276. 10.1023/a:1015299930849

Demb, J. B. (2007). Cellular Mechanisms for Direction Selectivity in the Retina. Neuron, 55(2), 179–186. 10.1016/j.neuron.2007.07.001

Dhande, O. S., Estevez, M. E., Quattrochi, L. E., El-Danaf, R. N., Nguyen, P. L., Berson, D. M., & Huberman, A. D. (2013). Genetic dissection of retinal inputs to brainstem nuclei controlling image stabilization. The Journal of Neuroscience: The Official Journal of the Society for Neuroscience, 33(45), 17797–17813. 10.1523/JNEUROSCI.2778-13.2013

Dhande, O. S., & Huberman, A. D. (2014). Retinal ganglion cell maps in the brain: Implications for visual processing. Current Opinion in Neurobiology, 24(1), 133–142. 10.1016/j.conb.2013.08.006

Douglas, R. M., Alam, N. M., Silver, B. D., McGill, T. J., Tschetter, W. W., & Prusky, G. T. (2005). Independent visual threshold measurements in the two eyes of freely moving rats and mice using a virtual-reality optokinetic system. Visual Neuroscience, 22(5), 677–684. 10.1017/S0952523805225166

Elstrott, J., Anishchenko, A., Greschner, M., Sher, A., Litke, A. M., Chichilnisky, E. J., & Feller, M. B. (2008). Direction selectivity in the retina is established independent of visual experience and cholinergic retinal waves. Neuron, 58(4), 499–506. 10.1016/j.neuron.2008.03.013

Feller, M. B. (1999). Spontaneous correlated activity in developing neural circuits. Neuron, 22(4), 653–656. 10.1016/s0896-6273(00)80724-2

Fried, S. I., Münch, T. A., & Werblin, F. S. (2002). Mechanisms and circuitry underlying directional selectivity in the retina. Nature, 420(6914), 411–414. 10.1038/nature01179

Giolli, R. A., Blanks, R. H. I., & Lui, F. (2006). The accessory optic system: Basic organization with an update on connectivity, neurochemistry, and function. Progress in Brain Research, 151, 407–440. 10.1016/S0079-6123(05)51013-6

Grubb, M. S., & Thompson, I. D. (2004). Visual response properties in the dorsal lateral geniculate nucleus of mice lacking the beta2 subunit of the nicotinic acetylcholine receptor. The Journal of Neuroscience: The Official Journal of the Society for Neuroscience, 24(39), 8459–8469. 10.1523/JNEUROSCI.1527-04.2004

Highstein, S. M., & Holstein, G. R. (2006). The anatomy of the vestibular nuclei. Progress in Brain Research, 151, 157–203. 10.1016/S0079-6123(05)51006-9

Horn, A. K. E., & Leigh, R. J. (2011). The anatomy and physiology of the ocular motor system. Handbook of Clinical Neurology, 102, 21–69. 10.1016/B978-0-444-52903-9.00008-X

Huang, L., Zhou, Y., Chen, W., Lin, P., Xie, Y., He, K., Zhang, S., Wu, Y., & Li, N. (2022). Correlations of FRMD7 gene mutations with ocular oscillations. Scientific Reports, 12(1), 9914. 10.1038/s41598-022-14144-7

Huberman, A. D., Feller, M. B., & Chapman, B. (2008). Mechanisms underlying development of visual maps and receptive fields. Annual Review of Neuroscience, 31, 479–509. 10.1146/annurev.neuro.31.060407.125533

Kay, J. N., De la Huerta, I., Kim, I.-J., Zhang, Y., Yamagata, M., Chu, M. W., Meister, M., & Sanes, J. R. (2011). Retinal ganglion cells with distinct directional preferences differ in molecular identity, structure, and central projections. The Journal of Neuroscience: The Official Journal of the Society for Neuroscience, 31(21), 7753–7762. 10.1523/JNEUROSCI.0907-11.2011

Kiraly, J. K., Harris, S. C., Al-Khindi, T., Dunn, F. A., & Kolodkin, A. L. (2024). PyOKR: A Semi-Automated Method for Quantifying Optokinetic Reflex Tracking Ability. Journal of Visualized Experiments: JoVE, (206), 10.3791/66779. 10.3791/66779

Mauss, A. S., Vlasits, A., Borst, A., & Feller, M. (2017). Visual Circuits for Direction Selectivity. Annual Review of Neuroscience, 40, 211–230. 10.1146/annurev-neuro-072116-031335

McLaughlin, T., Torborg, C. L., Feller, M. B., & O’Leary, D. D. M. (2003). Retinotopic map refinement requires spontaneous retinal waves during a brief critical period of development. Neuron, 40(6), 1147–1160. 10.1016/s0896-6273(03)00790-6

Medrea, I., & Cullen, K. E. (2013). Multisensory integration in early vestibular processing in mice: The encoding of passive vs. active motion. Journal of Neurophysiology, 110(12), 2704–2717. 10.1152/jn.01037.2012

Nalbach, H.-O., Thier, P., & Varjú, D. (1993). Binocular interaction in the optokinetic system of the crab Carcinus maenas (L.): Optokinetic gain modified by bilateral image flow. Visual Neuroscience, 10(5), 873–885. 10.1017/S0952523800006088

Oyster, C. W., & Barlow, H. B. (1967). Direction-selective units in rabbit retina: Distribution of preferred directions. *Science (New York*, N.Y.), 155(3764), 841–842. 10.1126/science.155.3764.841

Rasmussen, R. N., Matsumoto, A., Arvin, S., & Yonehara, K. (2021). Binocular integration of retinal motion information underlies optic flow processing by the cortex. Current Biology: CB, 31(6), 1165–1174.e6. 10.1016/j.cub.2020.12.034

Scudder, C. A., Kaneko, C. S., & Fuchs, A. F. (2002). The brainstem burst generator for saccadic eye movements: A modern synthesis. Experimental Brain Research, 142(4), 439–462. 10.1007/s00221-001-0912-9

Simpson, J. I. (1984). The accessory optic system. Annual Review of Neuroscience, 7, 13–41. 10.1146/annurev.ne.07.030184.000305

Simpson, J. I., Soodak, R. E., & Hess, R. (1979). The accessory optic system and its relation to the vestibulocerebellum. Progress in Brain Research, 50, 715–724. 10.1016/S0079-6123(08)60868-7

Stafford, B. K., Sher, A., Litke, A. M., & Feldheim, D. A. (2009). Spatial-Temporal Patterns of Retinal Waves Underlying Activity-Dependent Refinement of Retinofugal Projections. Neuron, 64(2), 200–212. 10.1016/j.neuron.2009.09.021

Stahl, J. S. (2004). Using eye movements to assess brain function in mice. Vision Research, 44(28), 3401–3410. 10.1016/j.visres.2004.09.011

Stewart, D. L., Chow, K. L., & Masland, R. H. (1971). Receptive-field characteristics of lateral geniculate neurons in the rabbit. Journal of Neurophysiology, 34(1), 139–147. 10.1152/jn.1971.34.1.139

Sugita, Y., Araki, F., Chaya, T., Kawano, K., Furukawa, T., & Miura, K. (2015). Role of the Mouse Retinal Photoreceptor Ribbon Synapse in Visual Motion Processing for Optokinetic Responses. PLOS ONE, 10(5), e0124132. 10.1371/journal.pone.0124132

Sun, C., Warland, D. K., Ballesteros, J. M., van der List, D., & Chalupa, L. M. (2008). Retinal waves in mice lacking the beta2 subunit of the nicotinic acetylcholine receptor. Proceedings of the National Academy of Sciences of the United States of America, 105(36), 13638–13643. 10.1073/pnas.0807178105

Sun, L. O., Brady, C. M., Cahill, H., Al-Khindi, T., Sakuta, H., Dhande, O. S., Noda, M., Huberman, A. D., Nathans, J., & Kolodkin, A. L. (2015). Functional Assembly of Accessory Optic System Circuitry Critical for Compensatory Eye Movements. Neuron, 86(4), 971–984. 10.1016/j.neuron.2015.03.064

Tabata, H., Shimizu, N., Wada, Y., Miura, K., & Kawano, K. (2010). Initiation of the optokinetic response (OKR) in mice. Journal of Vision, 10(1), 13. 10.1167/10.1.13

Tarpey, P., Thomas, S., Sarvananthan, N., Mallya, U., Lisgo, S., Talbot, C. J., Roberts, E. O., Awan, M., Surendran, M., McLean, R. J., Reinecke, R. D., Langmann, A., Lindner, S., Koch, M., Jain, S., Woodruff, G., Gale, R. P., Bastawrous, A., Degg, C.,…Gottlob, I. (2006). Mutations in FRMD7, a newly identified member of the FERM family, cause X-linked idiopathic congenital nystagmus. Nature Genetics, 38(11), 1242–1244. 10.1038/ng1893

Taylor, W. R., & Vaney, D. I. (2003). New directions in retinal research. Trends in Neurosciences, 26(7), 379–385. 10.1016/S0166-2236(03)00167-X

Thomas, M. G., Crosier, M., Lindsay, S., Kumar, A., Araki, M., Leroy, B. P., McLean, R. J., Sheth, V., Maconachie, G., Thomas, S., Moore, A. T., & Gottlob, I. (2014). Abnormal retinal development associated with FRMD7 mutations. Human Molecular Genetics, 23(15), 4086–4093. 10.1093/hmg/ddu122

Thurtell, M. J., & Leigh, R. J. (2011). Nystagmus and saccadic intrusions. Handbook of Clinical Neurology, 102, 333–378. 10.1016/B978-0-444-52903-9.00019-4

Tian, J., Zee, D. S., & Walker, M. F. (2007). Rotational and Translational Optokinetic Nystagmus Have Different Kinematics. Vision Research, 47(7), 1003–1010. 10.1016/j.visres.2006.12.011

Tiriac, A., Bistrong, K., Pitcher, M. N., Tworig, J. M., & Feller, M. B. (2022). The influence of spontaneous and visual activity on the development of direction selectivity maps in mouse retina. Cell Reports, 38(2), 110225. 10.1016/j.celrep.2021.110225

Trenholm, S., & Awatramani, G. B. (2015). Origins of spontaneous activity in the degenerating retina. Frontiers in Cellular Neuroscience, 9. 10.3389/fncel.2015.00277

Trenholm, S., Johnson, K., Li, X., Smith, R. G., & Awatramani, G. B. (2011). Parallel mechanisms encode direction in the retina. Neuron, 71(4), 683–694. 10.1016/j.neuron.2011.06.020

van Alphen, A. M., Stahl, J. S., & De Zeeuw, C. I. (2001). The dynamic characteristics of the mouse horizontal vestibulo-ocular and optokinetic response. Brain Research, 890(2), 296–305. 10.1016/s0006-8993(00)03180-2

Vaney, D. I., Peichl, L., Wässle, H., & Illing, R. B. (1981). Almost all ganglion cells in the rabbit retina project to the superior colliculus. Brain Research, 212(2), 447–453. 10.1016/0006-8993(81)90476-5

Wang, K., Hinz, J., Haikala, V., Reiff, D. F., & Arrenberg, A. B. (2019). Selective processing of all rotational and translational optic flow directions in the zebrafish pretectum and tectum. BMC Biology, 17(1), 29. 10.1186/s12915-019-0648-2

Watkins, R. J., Patil, R., Goult, B. T., Thomas, M. G., Gottlob, I., & Shackleton, S. (2013). A novel interaction between FRMD7 and CASK: Evidence for a causal role in idiopathic infantile nystagmus. Human Molecular Genetics, 22(10), 2105–2118. 10.1093/hmg/ddt060

Wei, W., & Feller, M. B. (2011). Organization and development of direction selective circuits in the retina. Trends in Neurosciences, 34(12), 638–645. 10.1016/j.tins.2011.08.002

Wibble, T., Pansell, T., Grillner, S., & Pérez-Fernández, J. (2022). Conserved subcortical processing in visuo-vestibular gaze control. Nature Communications, 13(1), 4699. 10.1038/s41467-022-32379-w

Wurtz, R. H. (2008). NEURONAL MECHANISMS OF VISUAL STABILITY. Vision Research, 48(20), 2070–2089. 10.1016/j.visres.2008.03.021

Wylie, D. R., & Frost, B. J. (1999). Responses of neurons in the nucleus of the basal optic root to translational and rotational flowfields. Journal of Neurophysiology, 81(1), 267–276. 10.1152/jn.1999.81.1.267

Yonehara, K., Balint, K., Noda, M., Nagel, G., Bamberg, E., & Roska, B. (2011). Spatially asymmetric reorganization of inhibition establishes a motion-sensitive circuit. Nature, 469(7330), 407–410. 10.1038/nature09711

Yonehara, K., Fiscella, M., Drinnenberg, A., Esposti, F., Trenholm, S., Krol, J., Franke, F., Scherf, B. G., Kusnyerik, A., Müller, J., Szabo, A., Jüttner, J., Cordoba, F., Reddy, A. P., Németh, J., Nagy, Z. Z., Munier, F., Hierlemann, A., & Roska, B. (2016). Congenital Nystagmus Gene FRMD7 Is Necessary for Establishing a Neuronal Circuit Asymmetry for Direction Selectivity. Neuron, 89(1), 177–193. 10.1016/j.neuron.2015.11.032

Yonehara, K., Ishikane, H., Sakuta, H., Shintani, T., Nakamura-Yonehara, K., Kamiji, N. L., Usui, S., & Noda, M. (2009). Identification of Retinal Ganglion Cells and Their Projections Involved in Central Transmission of Information about Upward and Downward Image Motion. PLOS ONE, 4(1), e4320. 10.1371/journal.pone.0004320

Yoshida, K., Watanabe, D., Ishikane, H., Tachibana, M., Pastan, I., & Nakanishi, S. (2001). A key role of starburst amacrine cells in originating retinal directional selectivity and optokinetic eye movement. Neuron, 30(3), 771–780. 10.1016/s0896-6273(01)00316-6

